# Ribosome biogenesis factor AtRRB1 confers pollen heat stress tolerance in Arabidopsis

**DOI:** 10.1101/2024.03.21.586164

**Authors:** Chunlian Jin, Cédric Schindfessel, Limin Sun, Burcu Nur Keçeli, Steffen Vanneste, Danny Geelen

**Affiliations:** Department of Plants and Crops, unit HortiCell, Faculty of Bioscience Engineering, Ghent University, Coupure Links 653, 9000 Ghent, Belgium

## Abstract

The plant male reproductive system is very sensitive to high temperature stress leading to a reduction in fertility. Damage caused by heat stress is restored by the activation of transcription and the synthesis of chaperones that regulate the heat stress response. Here we report that AtRRB1 is a homolog of the yeast ribosome chaperone protein Rrb1p. *AtRRB1* is an essential gene and a T-DNA insertion in the coding region impairs male and female gametogenesis. The heterozygous *rrb1-1* mutant shows decreased expression of AtRRB1 and increased transcription of the 60S ribosome proteins RPL3B and RPL4, in line with a chaperone role of AtRRB1 in ribosome biogenesis. Embryo sac development across ovules of a single pistil occurs uncoordinated and about half of the ovules abort. Half of *rrb1-1* pollen is substantially smaller and produce shorter pollen tubes than WT pollen. In contrast to the Col-0 pollen, smaller pollen is overly sensitivity to high temperature (24h at 32°C) treatment, specifically during the early bicellular microspore development stage. Heat stressed *rrb1-1* bicellular microspores accumulated excessively rough endoplasmic reticulum stacks, suggesting that loss of AtRRB1 activity causes an arrest in ER associated protein biosynthesis. These findings support a critical requirement for ribosome biogenesis and protein synthesis in bicellular microspores to recover from high temperature stress.

## Introduction

Ribosomes act as protein assembling robots either freely in the cytoplasm or attached to the rough endoplasmic reticulum (RER). They play an indispensable role in polypeptide chain formation, translating mRNA into proteins. The eukaryote ribosome is a large complex consisting of two parts, the 40s small subunit (SSU) and the 60s large subunit (LSU), consisting of multiple proteins and rRNA (Schmeing 2013). The main pathway of ribosome synthesis in eukaryotes is conserved. It starts in the nucleolus, where the components of pre-rRNA, 18S, 5.8S and 25S, are transcribed by Pol1 and undergo rRNA modification, forming the 90S pre-ribosome. The 90S pre-ribosome is then cleaved and matures into pre-40S and pre-60S. The final step is the maturation of 40S and 60S and their transport to the cytoplasm (Thomson et al. 2013). Eukaryote ribosome biogenesis involves ribosome proteins, snoRNPs (small nucleolar ribonucleoparticles) and hundreds of ribosome biogenesis factors (RBFs). Approximately 70% of the yeast RBFs are strongly conserved in Arabidopsis some of which partially complement the corresponding *S. cerevisiae* deletion/depletion strain, (*SG1-2^Lsg1p^*, *NOB1^Nob1p^*, *NUC1/2^Nsr1p^*, *NUG2^Nug2p^*, *PES^Nop7p^*, *REIL1^Rei1^*, *RRP44A^Rrp44/DIS3^* and *XRN3^Rat1p^*) while some cannot (*BRX1-1/1-2^Brx1p^*, *ENP1^Enp1p^*, *MTR4^Mtr4p^*, *NMD3^Nmd3p^*, *NOC4^Noc4p^*, *PWP2^Pwp2p^*, *RID2^Bud23^*, *RRP6-L2^Rrp6p^*, *RTL2^Rnt1p^* and *SWA2^Noc1p^*) (Weis et al. 2015). Loss of function Arabidopsis RBF mutants are affected in root and leaf development and epidermal patterning (Wieckowski and Schiefelbein, 2012; Petricka and Nelson, 2007; Abbasi et al., 2010; Hang et al., 2014; Weis et al., 2014). In addition, several RBFs have been shown to play a role in reproductive development. The Arabidopsis heterozygous mutants *pwp2^+/-^*, *rrp5^+/-^*, *enp1^+/-^* and *nob1^+/-^* produce siliques with aborted ovules, consistent with poor maternal transmission and an arrest of embryo development at the globular stage (Missbach et al. 2013). Pollen germination and pollen tube growth are impaired in *pwp2^+/-^*, *rrp5^+/-^*, *enp1^+/-^* and *nob1^+/-^* strongly reducing male transmission of mutant alleles (Missbach et al. 2013).

Yeast Rrb1p (Regulator of Ribosome Biogenesis1) is an essential RBF containing WD-repeats, a 44-60 amino acid sequence ending with Trp-Asp (WD), that is required for the synthesis and assembly of the 60S ribosome subunit and interacts with the ribosomal protein RPL3 (Schaper et al. 2001; Iouk et al. 2001). Disruption of *RRB1^YMR131C^* expression causes ribosome biogenesis defects and chromatin instability due to the inactivation of interaction with *YPH1^PES^* (*yeast pescadillo homolog 1*) and other members of the *YPH1^Pes^* complex, *RPL3*, *ERB1^BOP1^* and *ORC6* (Killian et al. 2004). Similarly, the depletion of GRWD1, the human homolog of RRB1, and the corresponding homologs of the PES complex, induce mitotic abnormalities (Killian et al. 2004). In plants, the PES complex is essential for biogenesis of the 60S ribosome large subunit (Cho et al. 2013). AtPES interacts with BOP1^ERB1^ and AtPEIP2^WDR12^ in the nucleolus and cofractionates with ribosome subunits (Zografidis et al. 2014). Depletion of either of these proteins leads to defects in cell division and primary root growth (Cho et al. 2013).

The homolog of *AtRRB1* in Arabidopsis was previously identified as *HEAT STRESS TOLERANT DWD 1* (*HTD1*; At2G19540), a Damaged DNA Binding1 (DDB1) binding WD40 protein and putative target of the ubiquitin E3 ligase Cullin4-RING (CRL4) (Kim et al. 2014). HTD1 transcription is increased upon heat shock (3h 37°C), congruent with a role in heat stress tolerance. Reduced expression of HTD1 in the promotor insertion mutant *htd1-1* (SALK_081295) improved regrowth after prolonged heat stress compared to the WT (Kim et al. 2014). In view of a putative role in thermotolerance we analysed the full knock out insertion mutant and discovered that it is embryo lethal and that heterozygous plants show enhanced expression of ribosomal proteins RPL3B and RPL4. We therefore renamed HTD1 to AtRRB1. The heterozygous *rrb1-1* mutant produces pollen of a normal size and a class of smaller pollen that under high temperature conditions accumulate rough endoplasmic reticulum stacks and eventually collapse. Our findings reveal a pivotal role for efficient protein synthesis in thermotolerance of pollen.

## Results

### AtRRB1 is an essential ribosome biogenesis factor

In search for candidate chaperones putatively involved in heat stress tolerance, we identified *AtRRB1,* the ortholog of yeast ribosome biogenesis factor *RRB1^YMR131C^* (Schaper et al. 2001; Iouk et al. 2001). *AtRRB1* (At2G19540) encodes a 469 amino acid protein consisting of six WD40 repeats and a chromatin assembly factor 1 domain (CAF1C-H4) at the N-terminal end of the protein and is conserved in yeast, plants, and humans (Figure S1). The mutant *rrb1-1* (GABI_837C04) has a T-DNA inserted in exon 11 at amino acid position 406 and *htd1-1* has a T-DNA insertion in the promotor region at position –297 (Figure 1A). Homozygous *rrb1-1* mutants were not recovered from selfed plants (Table S1), supporting its essential function in Arabidopsis, and heterozygous plants (hereafter *rrb1-1)* showed a reduction in *AtRRB1* expression in inflorescence tissue to about half of that in Col-0 (Figure 1B). The impaired *AtRRB1* transcription was concurrent with a significant increase in mRNA levels of RPL3B and RPL4 (Figure 1B), analogous to what has been reported for *RRB1^YMR131C^* mutants in yeast (Iouk et al. 2001). The expression of *RPL3A*, *RPL18*, and *RPL23AA* was unaltered, indicating that AtRRB1 depletion has a selective impact on the 60S ribosome biogenesis (Figure 1B). Also, in accordance with a putative chaperone function of AtRRB1 in the transport of ribosomal proteins from the cytoplasm to the nucleus (Pillet et al. 2015), AtRRB1-GFP localized in both compartments and was most abundant in the nucleolus (Figure 1C).

**Figure 1.**
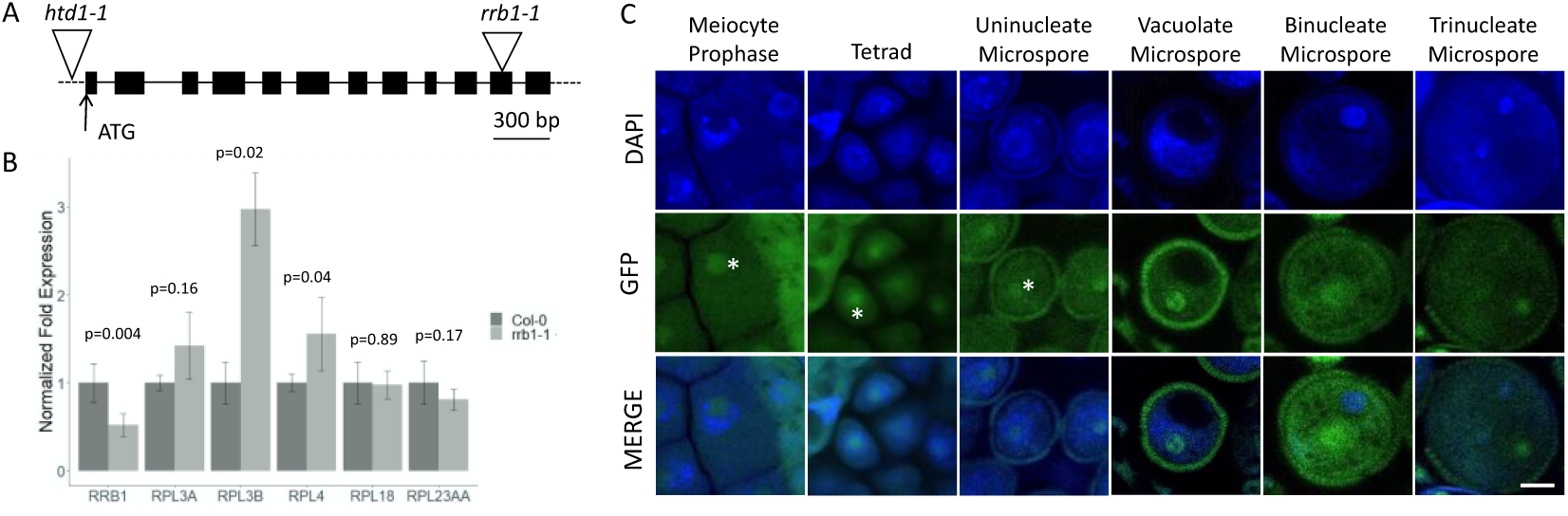
AtRRB1 is a putative ribosome biogenesis factor. A. Schematic structure of *AtRRB1* and the T-DNA insertion of *rrb1-1* in the 11^th^ exon and *htd1-1* in the promotor. B. Q-PCR of RRB1, RPL3A, RPL3B, RPL4, RPL18, and RPL23AA expression in inflorescence tissue from Col-0 and *rrb1-1* heterozygous plants. Normalized to the Col-0 control for every gene. Error bars represent standard errors. p-values are based on a linear model fit comparing Col-0 vs *rrb1-1* C. Localization of RRB1-GFP in developing microspores. RRB1-GFP accumulates primarily in the nucleolus (*) in early microspore development. Scale bar = 5µm.

### AtRRB1 has an essential function in reproduction

Although *rrb1-1* showed reduced *AtRRB1* and increased *RPL3B* and *RPL4* expression, these alterations did not have a clear impact on vegetative growth and development (Figure S2). However, *rrb1-1* plants are strongly reduced in fertility (Figure 2). Silique length and numbers of seeds per silique were significantly lower than in Col-0 and *htd1-1* (Figure 2B, C). Arabidopsis *rrb1-1* siliques contained 31±5 (n=29) seeds compared to 57±5 (n=29) in *htd1-1* and 58±5 (n=26) in Col-0. The reduced fertility phenotype of *rrb1-1* was fully complemented in lines transformed with p*R-RRB1*-GFP that express AtRRB1 fused to GFP from a promotor 315 bp upstream of the start codon (Figure 2B, C). Expression of *RRB1*-GFP from the LAT52 promotor (p*L-RRB1*-GFP) resulted in partial restoration of silique length and seed set (Figure 2A, B), indicating that the expression in mature pollen was not sufficient to fully restore the fertility defect.

**Figure 2.**
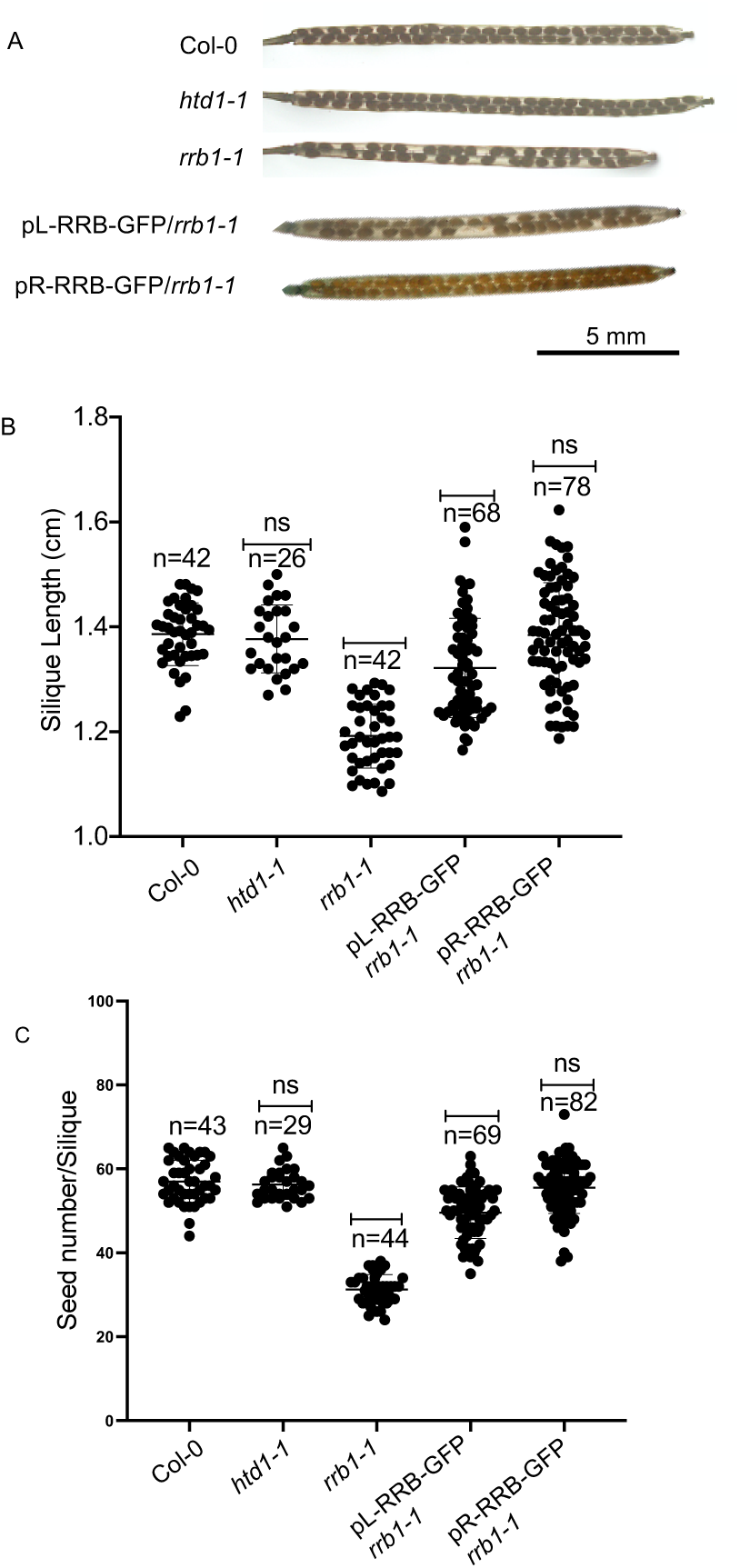
Reduced fertility of *rrb1-1*. A. Cleared siliques from Col-0, *htd1-1,* and *rrb1-1* and *rrb1-1* complemented with p*R-RRB1*-GFP and p*L-RRB1*-GFP. Scale bar= 5mm. B. Silique lengths of Col-0, *htd1-1,* and *rrb1-1* and *rrb1-1* complemented with p*R-RRB1*-GFP and p*L-RRB1*-GFP. Significantly shorter siliques were measured in *rrb1-1* and in *rrb1-1* complemented with p*L-RRB1*-GFP. C. Number of seeds per siliques. Significantly less seeds were found in *rrb1-1* and *rrb1-1* complemented with p*L-RRB1*-GFP.

### AtRRB1 is required for male and female gametophytic development

The absence of homozygous *rrb1-1* progeny indicated that the *rrb1-1* mutation is lethal, yet male and/or female transmission is partially functional generating heterozygous *rrb1-1*. The gametophytic transmission of the T-DNA insertion was determined by analyzing antibiotic resistance from reciprocal crosses of *rrb1-1* and Col-0. With *rrb1-1* as pollen donor about 25% of the progeny carried the T-DNA. The reciprocal cross generated about 18% *rrb1-1* progeny, indicating that transmission is slightly better via the male gametophyte (Table S2).

To investigate the defect in the female gametophyte, embryo sac development from ovules of individual pistils that were isolated from a single inflorescence were microscopically staged (Christensen et al. 1998). The Analysis 7 embryo sac stages were identified in about 4 to 5 pistils derived from subsequent flower buds starting with the most recently opened one (Figure 3; Table 1). Each pistil contained ovules at 2 to 3 consecutive developmental stages in Col-0 whereas in *rrb1-1* a single pistil contained up to 5 female gametophytic stages pointing to a reduced coordination of embryo sac development across the ovules. In separate analyses we observed that all ovules fully developed with synergids, egg cell, a central cell and 3 polar localized nuclei in *rrb1-1* (Figure 3A). After fertilization, half of the *rrb1-1* ovules developed embryos and the other half collapsed (Figure 3B).

**Figure 3.**
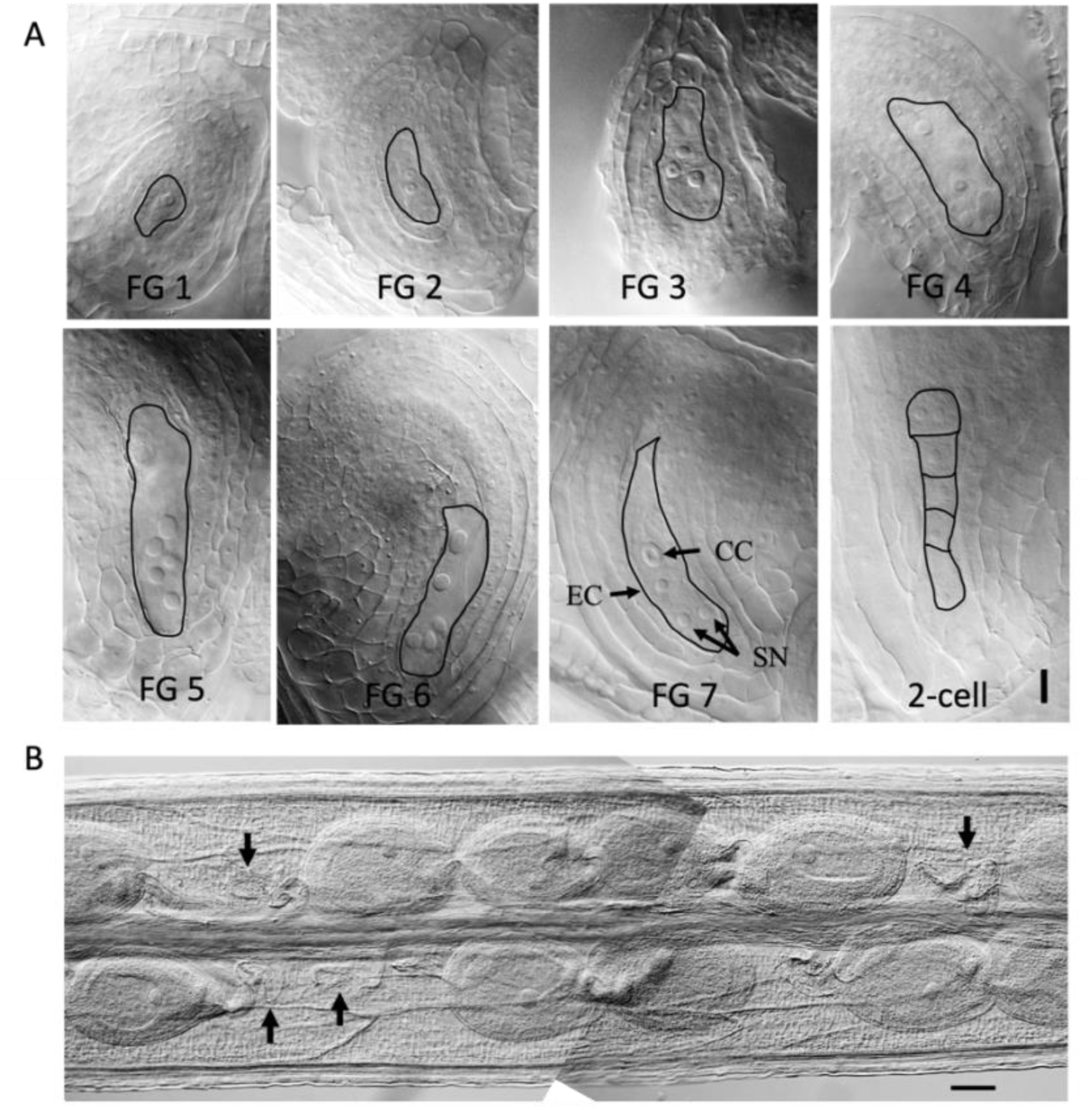
Megaspore mother cells development in *rrb1-1*. A. DIC imaging of ovules at stages FG1 to FG7 and an embryo at the 2-cell stage. EC egg; SN synergids; CC central cell. Scale bar= 10 μm. B. Subsection of a *rrb1-1* pistil with ovules carrying embryos at the globular stage at 3 days post fertilization. Collapsed ovules are indicated with an arrow.

**Table 1.** Frequency of female gametophyte stages found in Col-0 and *rrb1-1* pistils belonging to a single inflorescence. Staging was done in accordance with Christensen et al. (1998).

|  |  | FG1 | FG2 | FG3 | FG4 | FG5 | FG6 | FG7 |
| --- | --- | --- | --- | --- | --- | --- | --- | --- |
| Col-0 | pistil 1 | 17 | 8 |  |  |  |  |  |
|  | pistil 2 | 14 | 1 |  |  |  |  |  |
|  | pistil 3 | 14 | 11 | 1 |  |  |  |  |
|  | pistil 4 |  |  | 6 | 6 | 5 | 2 |  |
|  | pistil 5 |  |  |  |  | 4 | 6 | 4 |
|  | pistil 6 |  |  |  |  | 1 | 17 | 18 |
|  | pistil 7 |  |  |  |  | 3 | 23 | 19 |
|  | pistil 8 |  |  |  |  | 1 | 6 | 3 |
|  | pistil 9 |  |  |  |  |  | 11 | 34 |
|  | pistil 10 |  |  |  |  |  | 5 | 35 |
| *rrb1-1* | pistil 1 | 2 | 14 | 5 | 3 |  |  |  |
|  | pistil 2 | 5 | 6 | 6 | 1 | 2 | 1 |  |
|  | pistil 3 |  | 1 | 6 | 16 | 7 | 4 |  |
|  | pistil 4 |  |  | 1 | 10 | 9 | 12 | 9 |
|  | pistil 5 |  |  | 2 | 13 | 7 | 8 | 4 |
|  | pistil 6 |  |  | 1 | 0 | 4 | 6 | 29 |

To investigate pollen formation in *rrb1-1*, anthers were Alexander stained (Figure 4A). Col-0 and *rrb1-1* anthers were stacked with alive, pink stained pollen. Scanning electron microscopy identified two types: pollen of normal size and shape and pollen that were substantially smaller and slightly deformed (Figure 4B). Staining with DAPI showed that most *rrb1-1* pollen comprised a regular tricellular configuration and about 2% contained a single sperm that appeared larger than the sperm in the tricellular pollen (Figure 4C). In the *qrt^-/-^* mutant background, *rrb1-1* quartet structures contained 2 small and 2 normal pollen revealing the gametophytic phenotype (Figure 4D). Pollen size analysis showed that *rrb1-1* produced two populations of equal abundance: normal-sized pollen of ∼20.5 μm diameter and slightly smaller pollen of ∼17.7 μm diameter (Figure 4E). This phenotype was fully recovered in the complementing AtRRB1-GFP lines with either the AtRRB1 or LAT52 promotor (Figure 4E). Since both Col-0 and *rrb1-1* pollen appear viable (Figure 4A), pollen germination was considered as a possible cause for the poor transmission of the *rrb1-1* mutation. In vitro pollen germination assays showed that Col-0 and *rrb1-1* produced pollen tubes, confirming their viability (Figure S3). The tubes of *rrb1-1* pollen were however much shorter for about half of the pollen (Figure S3, 4F). While Col-0 pollen tube lengths were Gaussian distributed around the 165-185 μm maximum, *rrb1-1* pollen showed two peak maxima: one at 25-45 μm and a second around the size of the Col-0 pollen (Figure 4F). The short pollen tube length likely impairs the fertilization competence of *rrb1-1* pollen, causing a reduction in the transmission of the *rrb1-1* mutation.

**Figure 4.**
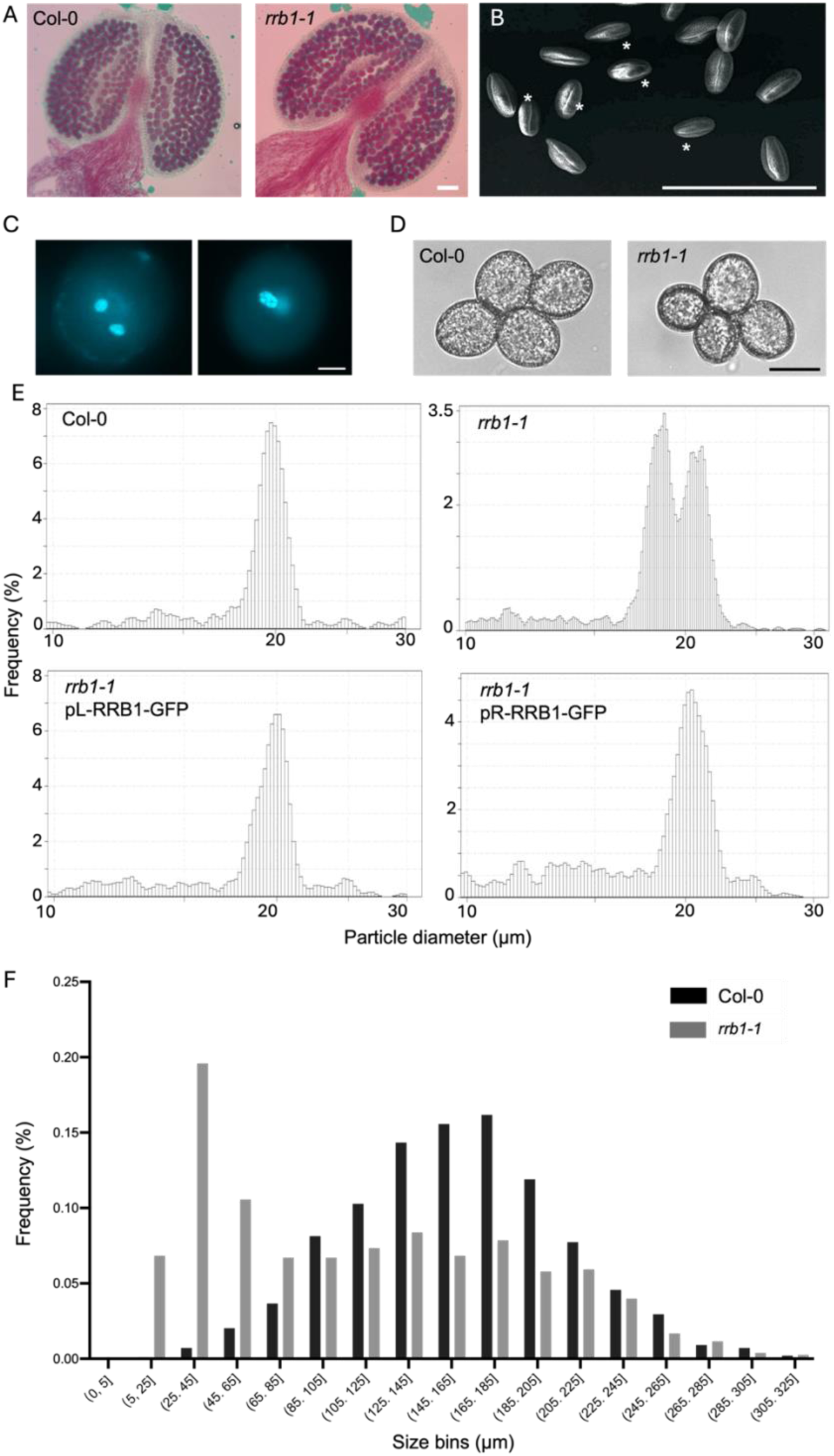
The *rrb1-1* pollen phenotype. A. Anthers stained with Alexander stain (size bar is 100 µm). A. SEM image of mature *rrb1-1* pollen with slight deformations and smaller pollen (*). C. DAPI stained rrb1-1 pollen, top normal tricellular configuration, bottom single sperm mature pollen (size bar is 4 µm). D. Pollen from *qrt*^-/-^ in the Col-0 or *rrb1-1* background. E. Pollen size distribution measured with a Coulter Counter. F. Distribution of pollen tube length of Col-0 (n=983) and *rrb1-1* (n=776) pollen germinated *in vitro*. The pollen tubes of Col-0 show a Gaussian distribution peaking at interval 165-185 μm. The pollen tubes from *rrb1-1* show 2 populations, a group with short tube around 25-45 μm and a second overlapping with those from Col-0.

### AtRRB1 confers pollen heat stress tolerance

To test whether AtRRB1 is involved in thermotolerance in pollen, we applied a 24h 32°C heat treatment that was previously shown to affect meiosis (De Storme and Geelen 2020). Pollen size was monitored for 5 consecutive days after heat (DAH). The heat treatment did not alter the size distribution of Col-0 and *htd1-1* pollen (Figure 5). However, the small pollen population of *rrb1-1* pollen (∼17.7 μm) collapsed at 2 DAH forming smaller particles (<15 μm). The small pollen population started to restore at 4 DAH (Figure 5). To verify whether specifically *rrb1-1* mutant pollen collapsed, emasculated Col-0 flowers were fertilized with pollen harvested at 3 DAH. Sulfadiazine selection of progeny showed that 22 seedlings out of 341 contained the *rrb1-1* T-DNA, a transmission rate of 6%, much lower than unstressed pollen (Table S2). To check what microspore development stage was heat sensitive, we staged every consecutive flower bud in an inflorescence by dissecting and DAPI staining a single anther before the heat treatment. Following heat treatment, pollen particle size analysis of the remaining anthers was performed as the flowers opened during the 5 consecutive days after the heat treatment. This revealed that the 18µm *rrb1-1* pollen population was eliminated upon the heat treatment during the bi-nucleate stage (Figure 5).

**Figure 5.**
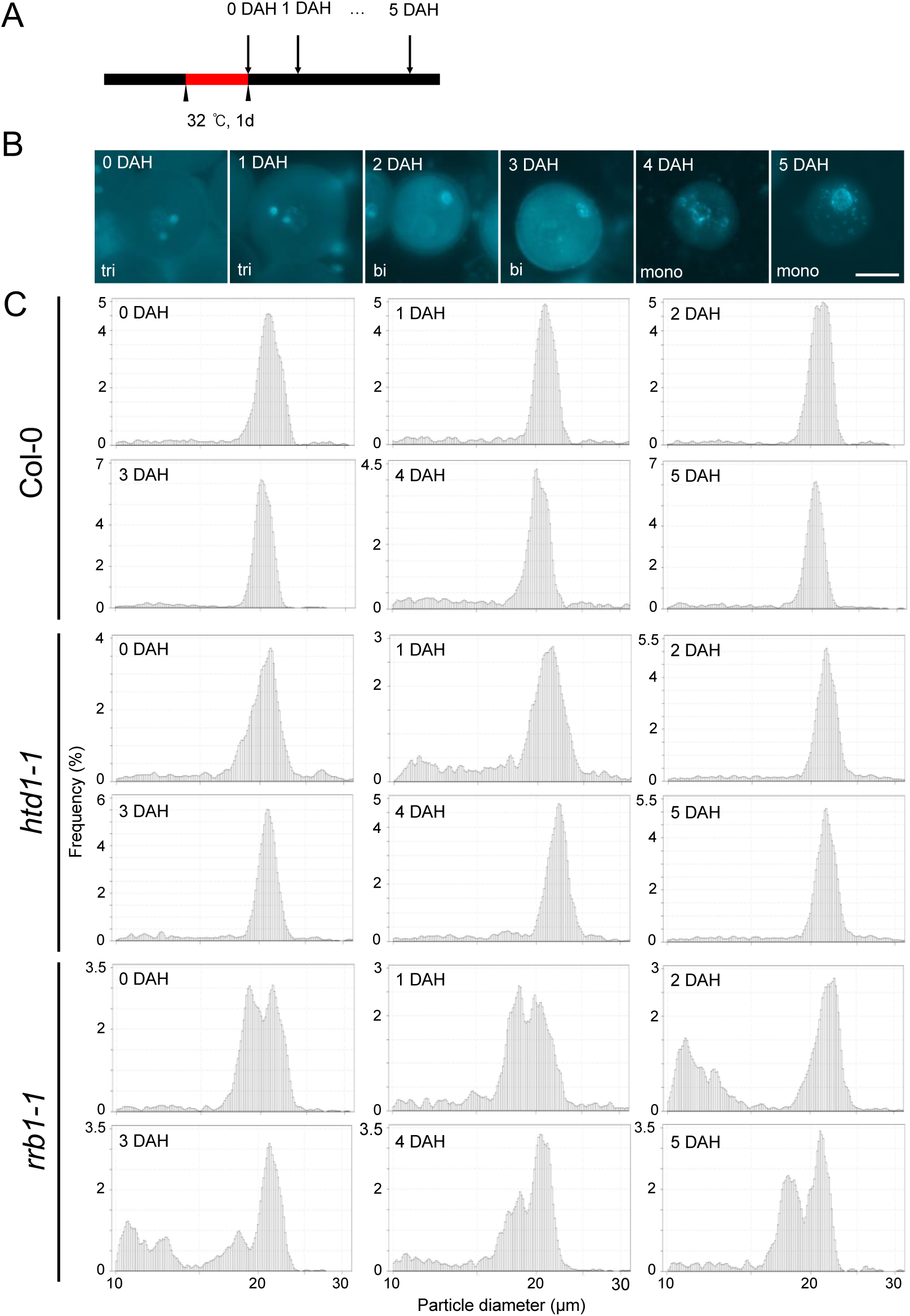
Pollen particle size distributions of Col-0, *rrb1-1*, and *htd1-1.* A. Pollen were collected 0-5 days after 24h heat treatment at 32°C (DAH). B. DAPI staining of pollen harvested at 0-5 DAH. Coulter counter size distribution histograms of Col-0, *rrb1-1* and *htd1-1* pollen isolated from consecutive days after the heat treatment.

### Polysomic ER stacks accumulate in heat stressed ***rrb1-1*** pollen

To gain insight into cellular defects occurring in heat stressed pollen, pollen were imaged at the ultrastructural level. Thin sections of Col-0 and *rrb1-1* pollen cultivated at 21°C showed an organized cytoplasm with a centrally located vegetative nucleus and two sperm cells, mitochondria, Golgi, and dense and transparent vesicles (Figure 6). In about half of the *rrb1-1* pollen, the cytoplasm contained irregularly clustered stacks of ER sheets which in Col-0 pollen consisted of less abundant ER stacks (Figure 6). Heat stress applied 24h before anthesis to *rrb1-1* caused a massive accumulation of ER stacks (> 2 μm in diameter) (Figure 6). The *rrb1-1* ER sheets were decorated with electrodense dots (rough ER) indicating that assembled ribosomes were targeted to these ER stack (Figure 6). Flowers that were heat stressed 48h before anthesis produced next to collapsed pollen spores in which transparent vesicles were engulfed by a double membrane. Heat stressed Col-0 pollen did not show excessive ER stacking nor engulfed vesicles.

**Figure 6.**
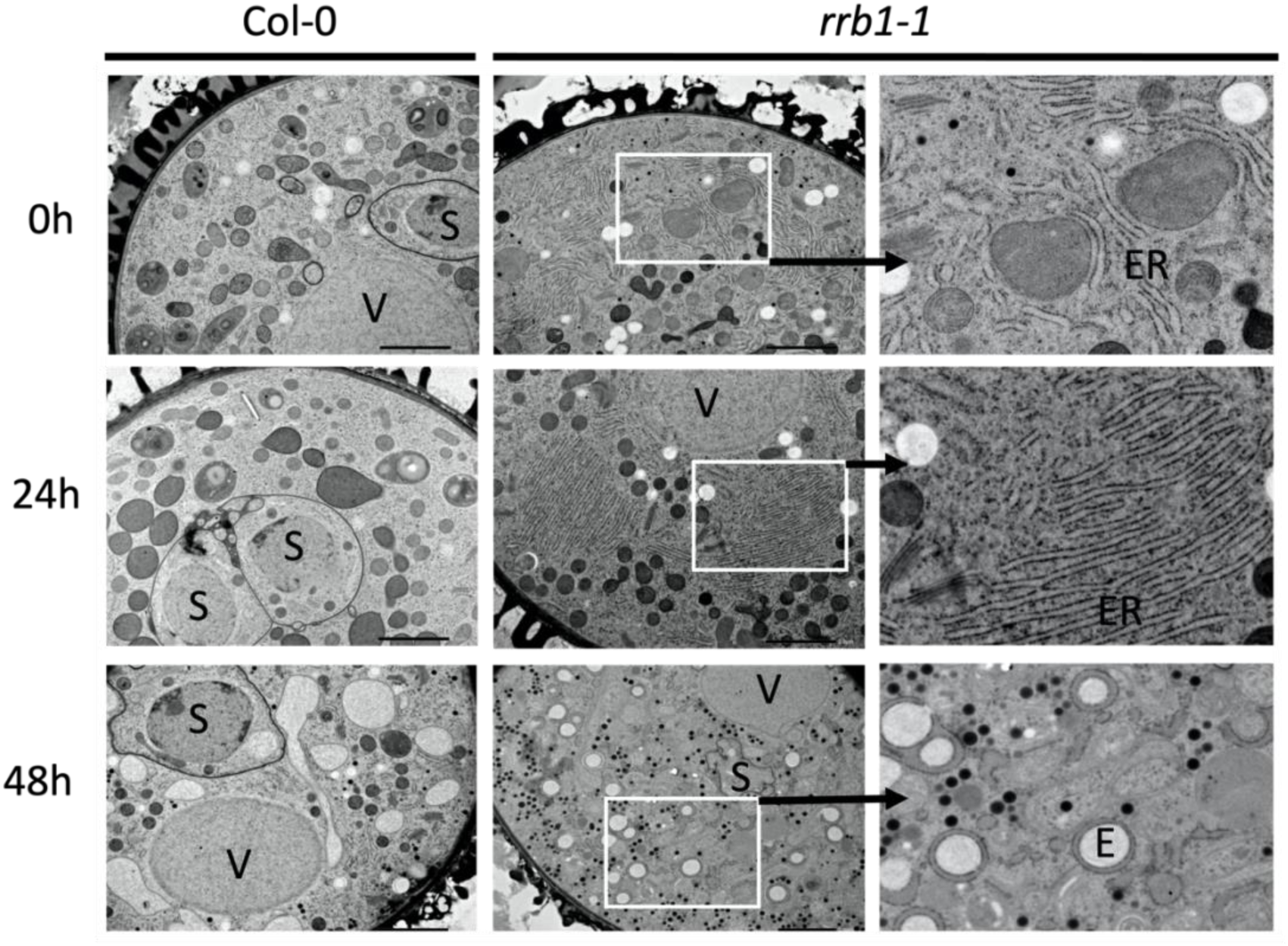
TEM of Col-0 and *rrb1-1* pollen after heat stress. Images were taken from pollen harvested at 0h, 24h and 48h after a 24h 32°C treatment. White frames are insets of enlarged images in the right panel. V, vegetative nucleus; S, Sperm cell; E, engulfed vesicle; ER, endoplasmic reticulum. Scale bar = 2μm.

## Discussion

### AtRRB1 is a ribosome biogenesis factor required for male and female gametogenesis

The *AtRRB1/HTD1* gene was identified as a homolog of the yeast *RRB1^YMR131C^* ribosome biogenesis factor essential for cell viability and assembly of the 60S ribosomal subunit (Schaper et al. 2001). Like in yeast, *AtRRB1* is essential for survival, preventing the isolation of homozygous *rrb1-1* lines. The *rrb1-1* heterozygous mutant shows reduced *AtRRB1* and increased *RPL3B* and *RPL4* mRNA levels respectively. Yeast RPL3 and RPL4 protein and mRNA levels are tightly controlled by their respective chaperones Rrb1p and Acl4p ensuring that just enough ribosomal protein is produced as the cell requires (Pillet et al. 2022). Conservation of a similar feedback regulation by AtRRB1 would explain why the heterozygous *rrb1-1* mutant displays no growth phenotype. Half of the male and female *rrb1-1* gametes show growth and cell division defects reminiscent to the gametophytic defects reported for several putative ribosome biogenesis factors (Shi et al. 2005; Li et al. 2009, 2010, 2019; Missbach et al. 2013; Cho et al. 2013; Ahn et al. 2016; Hao et al. 2017; Xiong et al. 2020). Expression of AtRRB1-GFP complements the reduced fertility of *rrb1-1* and reveals its cellular localization in the nucleolus, an important hub for ribosome assembly (Sáez-Vásquez and Delseny 2019). The molecular structure and phenotypic characteristics of the *rrb1-1* mutant strongly support that AtRRB1 is an ortholog of yeast Rrb1p and functions in ribosome assembly (Iouk et al. 2001).

AtRRB1/HTD1 is a member of the large WD40 protein family and belongs to a subclass that interacts with the CULLIN4 RING E3 ligase (CRL4) complex involved in protein ubiquitination and subsequent 26S proteolytic degradation (Lee et al., 2008; Kim et al. 2014). The CUL4 scaffold protein binds the RING finger protein DAMAGED DNA BINDING PROTEIN1 (DDB1) that recruits chromatin remodelling factors and chaperones that have a characteristic DWD motif (Fonseca and Rubio 2019). This suggests that AtRRB1 acts as a substrate receptor for proteins to be targeted for degradation. Putative candidates for regulation by CRL4 degradation are the putative AtRRB1 clients RPL3B and RPL4, or other ribosome assembly factors like PWP1 that was shown to be a target of CRL4 in animals (Han et al., 2020). AtRRB1 weakly interacts with HSP90-1 which may be important under conditions of heat stress to regulate the recruiting capacity of AtRRB1 (Kim et al., 2014).

Several other WD40 motif containing proteins have been associated with ribosome biogenesis: SLOW WALKER1 (SWA1) a homolog of yeast Noc1 with six WD40 repeats (Shi et al. 2005); PERIODIC TRYPTOPHAN PROTEIN 2 (PWP2), a homolog of yeast Pwp2 (Missbach et al. 2013), and YAO, with seven WD repeats (Li et al. 2010). The SWA1, PWP2 and YAO proteins are involved in nucleolar processing of pre-18S ribosomal RNA for 40S subunit maturation and play critical roles in fertility and female gametogenesis (Shi et al. 2005; Li et al. 2010; Missbach et al. 2013). While maternal transmission is strongly reduced in *rrb1-1*, it is totally blocked in *swa1, pwp2* and *yao* mutants, suggesting that these factors may have additional functions during embryo sac development or embryogenesis. This is further supported by a similar but less severe delay in embryo sac development observed in *rrb1-1* pistils compared to swa, pwp2 and yao mutants (Shi et al. 2005; Li et al. 2010; Missbach et al. 2013). The *rrb1-1* embryo sacs in mature pistils displayed an 8-celled polygonum type of configuration, indicating that the three gametophytic cell divisions were completed. Despite the full execution of the gametophyte program, only half of the *rrb1-1* ovules developed into seeds. The other half of the fertilized ovules aborted around 3 days post fertilization when normal developing ovules was at the early/mid globular stage. We did not observe cell division defects at earlier *rrb1-1* ovule development stages in DIC microscopy that would reveal the cellular defect leading to ovule abortion.

AtRRB1 contributes to regular pollen development whereby half of the *rrb1-1* pollen was smaller in size, a phenotype that was also recorded for mutations in the RBFs NOB1 and ENP1 (Missbach et al., 2013). With the germination and viability of *rrb1-1* pollen being similar to that of WT, we assume that reduced transmission of the AtRRB1 T-DNA insertion is more tightly linked to poor pollen tube growth. Impaired pollen tube elongation has also been observed in the Arabidopsis mutant *snail1* that carries a T-DNA insertion in SNAIL1, an ortholog of the yeast RBF Ssf1 (Hao et al. 2017). The initiation and growth of the pollen tube is in many pollen species largely independent of transcription but critically dependent on translation of stored mRNAs (Honys et al. 2009). Large ribonucleoprotein particles containing silent mRNA and ribosomal subunits are formed in the immature microspore that become activated upon pollen germination allowing a rapid translation of the temporally stored mRNA. Protein profiling of in vivo-grown pollen tubes revealed abundant expression of ribosome biogenesis factors underlining the critical requirement of a performant translation machinery during pollen tube growth (Lin et al. 2014). Hence, it is likely that the aberrant tube growth of part of the *rrb1-1* pollen is caused by an impaired assembly of functional ribosome complexes.

### Heat oversensitivity of rrb1 pollen

Damage to the reproductive system inflicted by heat is highly complex, whereby defects in male meiosis, microspores development, premature degradation of the tapetum, and pollen tube growth all contribute to reduced pollen viability and fertility (Giorno et al. 2013; De Storme and Geelen 2014; Chaturvedi et al. 2021). Moreover, the severity of the damage and the type of tissue affected depends on the temperature threshold, the amplitude and ramp of the temperature shift, the duration of the heat stress, and the historical context that may involve an eventual previous exposure to high temperature leading to the so-called priming response (Jagadish et al. 2021). The broad spectrum of defects induced by high temperature complicates the molecular analysis of underlying processes and so far, very few molecular factors have been identified that contribute to heat stress tolerance of pollen. In rice, the *GROWTH REGULATING FACTOR 4* (*OsGRF4*), and *HEAT SHOCK FACTOR60-3B* (*OsHSP60-3B*) are required to secure viability and seed-setting rate under heat stress (Mo et al. 2023; Lin et al. 2023). And in maize, improved tolerance of meiosis to heat stress has been reported in lines overexpressing *HEAT SHOCK PROTEIN101* (*HSP101*) (Li et al. 2022). Here we treated flowering Arabidopsis with a modest heat stress (32°C) over an extended period (24 h) and uncover the importance of intact ribosome biogenesis as a heat stress tolerance mechanism for pollen viability.

The small-sized pollen fraction in *rrb1-1* is hypersensitive to high temperature. Considering that AtRRB1 levels will be low in this pollen, the strong sensitivity to heat stress is in apparent conflict with the increased heat tolerance of *htd1-1* seedlings that is impaired in expression of AtRRB1 (Kim et al. 2014). However, *htd1-1* does not produce small pollen, a phenomenon that was also reported for several other ribosome biogenesis factors (Missbach et al. 2013). We therefore assume that *htd1-1* does not suffer from insufficient ribosome biogenesis and protein synthesis during microsporogenesis.

### Protein translation as important machinery to overcome heat stress damage

Studies investigating the impact of heat stress on vegetative tissue have unveiled the heat shock response that involves several conserved heat shock transcription factors (HSR) that move from the cytosol to the nucleus and upregulate the expression of heat shock proteins (HSP) that encode chaperone proteins involved in maintaining the activity of vital cytosolic proteins (Bourgine and Guihur, 2021). Heat stress also affects membrane associated proteins and those residing in the lumen of ER, Golgi and within the endosome. The accumulation of unfolded proteins causes ER stress that activates the unfolding response (UPR) pathway conserved in eucaryotes. Steady state activity of the UPR signalling pathway genes is critical for regular pollen development and male sterility (Singh et al. 2021). The molecular signature of the UPR is the upregulation of genes encoding endoplasmic reticulum (ER) chaperones and components of the ER-associated degradation (ERAD) system (Martínez and Chrispeels 2003). The UPR upregulates membrane-associated transcription factors, such as bZIP17 and –28, and on the other hand the splicing of bZIP60 messenger RNA by INOSITOL REQUIRING ENZYME 1 (IRE1) (Nagashima et al. 2011; Deng et al. 2011). Arabidopsis mutants that are UPR-deficient (lacking bzip28, bzip60, bZIP17, or IRE) are sensitive to heat stress and show a more pronounced reduction in fertility (Deng et al. 2016; Zhang et al. 2017; Gao et al. 2022). Maintaining protein homeostasis is therefore critical for pollen viability under heat stress (Singh et al. 2021). However, it remains unknown how the overaccumulation of misfolded protein in ER leads to pollen collapse (Rieu et al., 2017).

In a recent study, five Immune-associated nucleotide-binding protein (IAN) genes (IAN2 to IAN6) were associated with variation in heat tolerance at the reproductive stage in Arabidopsis thaliana accessions (Lu et al. 2021). The loss of function of IAN genes enhances the expression of HSR and UPR genes and reduces heat induced cell death by suppressing Bcl-2-associated athanogene 7 (BAG7), an ER-localized UPR protein that inhibits cell death (Williams et al. 2010). The IAN2 to IAN6 proteins are partially localized to the ER and provide a possible link between heat induced ER stress and the activation of cell death.

The double *ire1a ire1b* double knockout mutant is fertile at room temperature, but male sterile at slightly elevated temperatures (Deng et al. 2016). The reduced viability of mature pollen was associated with altered pollen coat composition and highly vacuolated tapetal cells at an early pollen stage (Deng et al. 2016). Because *rrb1-1* lines are heterozygous microspore surrounding tissue is not affected and unlikely responsible for the collapse of the high temperature treated *rrb1-1* pollen. Instead, we found that the pollen produced super numerous rough ER cisternae that are likely coated with dysfunctional ribosomes stalled at the ER surface. At a later stage the *rrb1-1* pollen produced membrane structures engulfing endosome vesicles and other vesicles, suggestive for activation of autophagy, a process that associated with and promotes cell death (Feng et al. 2022). We therefore speculate that, under normal conditions, mutant pollen carries sufficient AtRRB1 protein for limited ribosome biogenesis and keep ER homeostasis by accumulating ER stacks to maintain viable albeit smaller pollen. Under heat stress, WT pollen survives through the activation of multiple heat response pathways whereas mutant *rrb1-1* pollen cannot fulfil the demand for protein synthesis and overloads the ER with misfolded proteins, activating autophagy and pollen cell death.

## Materials and Methods

### Plant material and growth conditions

Arabidopsis ecotype Columbia-0 (Col-0), *qrt^-/-^,* T-DNA insertion lines GABI_837C04 and Sail_37_E12 were ordered from Nottingham Arabidopsis Stock Center (NASC). Seeds were grown on K1 medium [2.514 g/L MS (without vitamins, Labconsult^®^), 10 g/L sucrose (Tienen-Tirlemont^®^), 100 mg/L myo-inositol (Duchefa^®^) and 0.5 g/L MES (Duchefa^®^), 8 g/L Plant Tissue Culture agar (International Medical^®^)] for 7 days (21°C, 16h/8h day/night) after 2-days vernalization at 4°C in the dark. Transgenic lines were selected on K1 medium with 75 mg/l sulfadiazine and further cultured on Jiffy^®^ substrate in a growth room with 200 μmol/m^2^/s fluorescent light 16h/8h day/night and 70% RH. Plants were genotyped by PCR with primers FP: 5’-GTCCAGCCGAAGAAAACGTG-3’; RP: 5’-TCAGACGGAAGCGTGTTCTG-3’ O8409: 5’-ATATTGACCATCATACTCATTGC-3’. For heat treatments, plants were transferred to a climate chamber (Panasonic MLR-352H-PE) at 32°C for 24 or 48 hours as indicated. The photoperiod and relative humidity were the same as the control growth condition.

### Real-time quantitative PCR

Total RNA was extracted from flower buds using the Promega ReliaPrep RNA Tissue Miniprep System. Following by reverse transcription to cDNA using the GoScript Reverse Transcription System (Promega, A5001). Real-time quantitative PCR (RT-qPCR) was performed using the GoTaq QPCR Master Mix (Promega, A6001) according to the manufacturer’s instructions. EF1αA4 was used as the internal control. The primers used were: RRB1F, CAGAGTTCAATGCACAGACCAAGG; RRB1R, GCCAGTGAAGTTCCTTCAGATC; RPL3AF, TTGGTGCGTGGCATCCTGCT; RPL3AR, TGGCTGTGTGTGCCTCAGTACCA; RPL3BF, CTCATCGTGGTCTTCGCAAG; RPL3BR, TGAGTCTCCTGGCCAACTTT; RPL4F, GGTTCCAAGAGAGCGGTTC; RPL4R, GCCTCCTCCTTAGTGACGAC; GCCAGTGAAGTTCCTTCAGATC; RPL18F, GAGGTGAATGCTTAACCTTTGACC; RPL18R, AGGTCCGAAATGCTTCACTGC; RPL23AAF, GAAGATGTATGACATCCAGACC; RPL23AAR, TCTGGTGTAAGCCTCACGTAAGCC; EF1αA4F, AGGCTGGTATCTCTAAGGATGGTCA; EF1αA4R, GGATTTTGTCAGGGTTGTATCCG.

### Microscopy

Siliques were dissected under a binocular Olympus SZX9 microscope and imaged with an Olympus DP21 camera. Immature ovules were cleared with chloral hydrate and imaged with differential interference contrast (DIC) light microscopy as described by Franks (2016). Pollen viability was determined using Alexander staining and the nuclear organization was visualized using DAPI staining (De Storme and Geelen, 2011). Pollen were germinated in vitro and the tube length determined using Image J (Li, 2011). Images were taken using Olympus IX81 inverted fluorescence microscope equipped with an X-Cite Series 120Q UV lamp and an Olympus XM10 camera. For the visualization of AtRRB1-GFP the whole-mount anther DAPI staining protocol of Capitao et al. (2021) was followed. Confocal microscopy was performed on a Nikon A1R HD25.

### Pollen size analysis

Pollen volumetric analysis was performed as previously described (De Storme et al., 2013). Pollen were extracted in Isoton II^®^ from 5∼10 mature flowers unless otherwise stated. Flower tissue was removed, and the suspension analyzed using a Coulter counter Multisizer 3^®^.

### Electron and scanning microscopy

Air-dried pollen was sputter-coated with gold and imaged in a Hitachi S-3000N variable pressure scanning electron microscope. For transmission electron microscopy anthers from mature flowers were fixed 1h in 4% paraformaldehyde + 2.5% glutaraldehyde in 0.1 M Cacodylate at room temperature, pH7.2 under vacuum and further incubated at 4℃ overnight in fresh fixative. Samples were 3 times washed in 0.1 M Cacodylate buffer for 30 min, rotating at 4℃ and then post fixed in reduced 1% OsO_4_ (1 ml 4% OsO_4_, 3 ml 0.134M NaCaco (pH=7.4), 66mg K_3_Fe(CN)_6_) overnight at 4℃, rotating. Then samples were washed 3 times in ddH_2_O, rotating at 4℃, 30 min and stained in 1% uranyl acetate for 1h in the dark at 4℃. Samples were dehydrated at 4℃ in 8 steps in increasing concentrations ethanal, infiltrated with propylene oxide followed by embedding in Spurr resin. Semi thin sections were made with an ultra-microtome (Leica EM UC6) at 0.5 μm and stained with 1% toluidine blue and 2% borax in distilled water. Ultrathin sections of gold interference color were made with an ultra-microtome and post-stained in a Leica EM AC20 for 40 min in uranyl acetate at 20℃ and for 10 min in lead citrate. Sections were collected on formvar-coated copper slot grids. Grids were imaged with a JEM 1400 plus TEM.

## Acknowledgements

We are grateful for Christophe Petit, Ellen Van Gysegem and Patricia Delaere who provided technical and logistical support. We thank Riet De Rycke for support with the electron microscopy sample preparation and imaging.

## Author contribution

CJ, CS, LS, BNK performed experiments. CJ, CS, LS and DG prepared the figures. DG developed the concept and wrote first draft. DG and SV edited the manuscript.

