## Supplemental figures and tables for "Ribosome biogenesis factor AtRRB1 confers pollen heat stress tolerance in Arabidopsis"

### Supplementary figures

### Figure S1

```
AtRRB1          ------------------------------------------------------------
Os11g0134500    ------------------------------------------------------------
GRWD1           ------------------------------------------------------------
Rrb1p           MSKRSIEVNEEQDRVVSAKTESHSVPAIPASEEQDAPMNDLEEQLSDEFDSDGEIIEIDG 60

AtRRB1          -----MGRNVKTKAKRKNKKKAEASSSEIPSIPTRVWQPG-VDTLEDGEELQCDPSAYNS 54
Os11g0134500    -----MGRKIKVKKKKASSKKAEASSSRVPSAPAKVWQPG-VDTLEEGEELQFDPQAYNY 54
GRWD1           ----MAARKGRRRTCETGEPMEAESGDTSSEGPAQVYLPGRGPPLREGEELVMDEEAYVL 56
Rrb1p           DDEINDEDDLRKKQEEAETLVQKDQSEGNKEKIQELYLPHMSRPLGPDEVLEADPTVYEM 120
                        . : :  .         ...   .   .:: *    .*  .* *  *  .*

AtRRB1          LHGFHVGWPCLSFDILGDKLGLNRTEFPHTLYMVAGTQAEKAAHNSIGLFKITNVSGKRR 114
Os11g0134500    LRGFNIGWPCLSFDVVRDQLGLVRSEFPHTLYGVAGTQAERATWNYIGIFKICNINGKKR 114
GRWD1           YHRAQTGAPCLSFDIVRDHLGDNRTELPLTLYLCAGTQAESAQSNRLMMLRMHNLHGTKP 116
Rrb1p           LHNVNMPWPCLTLDVIPDTLGSERRNYPQSILLTTATQSSRKKENELMVLALSNLAKTLL 180
                 :  :   ***::*:: * **   * : * ::   :.**:.    * : :: : *:  .

AtRRB1          DVVPKTFGNGEDEDEDDEDDSDSDDDDGDEASKTPNIQVRRVAHHGCVNRIRAMPQ-NSH 173
Os11g0134500    EPIPASAIDGDSD--MDSESSSDEEDEAVNEDTMPILHLKKVAHAGCVNRIRSMNQ-EPH 171
GRWD1           PPS-----------EGSDEEEEEDEEDEEERKPQLELAMVPHYGGINRVRVSWLGEEP 164
Rrb1p           KDDN----------EGEDDEEDDEDDVDPVIENENIPLRDTTNRLKVSPFAISNQ--EV 227
                          ..:....::*:         : :  ..:   :. .

AtRRB1          ICVSWADSGHVQVWDMSSHLNALAESETEGKDGTSPVLNQAPLVNFSGHK-DEGYAIDWS 232
Os11g0134500    ICATWGDTGHVQVWDFSSFLNSLAESGAVAHNEDDRIHNHVPVKIFGSHK-DEGYAIDWS 230
GRWD1           VAGVWSEKGQVEVFALRRLLQVVEEPQALAAFLRDEQAQMKPIFSFAGHM-GEGFALDWS 223
Rrb1p           LTATMSENGDVYIYDLAPQSKAFSTPGYQIPKSVKR-----PIHTVKNHGNVEGYGLDWS 282
                :    .:.*.* :: :    : .  .         .       *:  . .*   **:.:***

AtRRB1          PATAG-RLLSGDCKSMIHLWEPASG-SWAVDPIPFAG-HTASVEDLQWSPAEENVFASCS 289
Os11g0134500    PLVTG-RLVSGDCNKCIHLWEPTSN-SWNVDTNPFVG-HTASVEDLQWSPTEADIFASCS 287
GRWD1           PRVTG-RLLTGDCQKNIHLWTPTDGGSWHVDQRPFVG-HTRSVEDLQWSPTENTVFASCS 281
Rrb1p           PLIKTGALLSGDCSGQIYFTQRHTS-RWVTDKQPFTVSNNKSIEDIQWSRTESTVFATAG 341
                *      *::***.  *::      .  * .*  **.  :. *:**:*** :*  :**:..

AtRRB1          VDGSVAVWDIRLG---KSPALSFKAHNADVNVISWNRLASCMLASGSDDGTFSIRDLRLI 346
Os11g0134500    ADRTISIWDIRTG---KKPCISVRAHNADVNVISWNRLASCMIASGCDDGSFSIRDLRLI 344
GRWD1           ADASIRIWDIRAAPSKACMLTTATAHDGDVNVISWSRREPFLLSGG-DDGALKIWDLRQF 340
Rrb1p           CDGYIRIWDTRSK--KHKPAISVKASNTDVNVISWSDKIGYLLASGDDNGTWGVWDLRQF 399
                 *  : :** *           :  * : *******.     :::.* *:*:  : *** :

AtRRB1          KGG----DAVVAHFEYHKHPITSIEWSAHEASTLAVTSGDNQLTIWDLSLEKDEEEEAEF 402
Os11g0134500    KD-----DSLVAHFEYHKHPITSVEWSPHEPSTLAVSSADHQLTIWDLSLEKDAEEEAEF 399
GRWD1           KS-----GSPVATFKQHVAPVTSVEWHPQDSGVFAASGADHQITQWDLAVERDPE----- 390
Rrb1p           TPSNADAVQPVAQYDFHKGAITSIAFNPLDESIVAVGSEDNTVTLWDLSVEADDEE---- 455
                .          ** :. *  .:**: : . : . .*. . *: :* ***::* * *

AtRRB1          NAQTKELVNTPQDLPPQLLFVHQGQKDLKELHWHNQIPGMIISTAGDGFNILMPYNIQNT 462
Os11g0134500    RARMREQADAPEDLPPQLLFVHQGQKDLKELHWHPQIPSMIISTAADGFNMLMPSNIDTT 459
GRWD1           -AGDVEADPGLADLPQQLLFVHQGETELKELHWHPQCPGLLVSTALSGFTIFRTISV--- 446
Rrb1p           IKQQAAETKELQEIPPQLLFVH-WQKEVKDVKWHKQIPGCLVSTGTDGLNVWKTISV--- 511
                                ::* ******  :.::*:::** * *. ::**. .*:.:  . .:

AtRRB1          LPSELPA 469
Os11g0134500    IREADA- 465
GRWD1           -------
Rrb1p           -------
```

Figure 1S. CLUSTAL 2.1 amino acid alignment of RRB1 like sequences from yeast (Rrb1p), human (GRWD1) and rice (Os11g0134500).
RRB orthologs of yeast (Rrb1p), human (GRWD1), rice (Os11g0134500) and Arabidopsis (AtRRB1) are aligned. The DWD motif is highlighted in grey and the CAF1C-H4 binding domain is illustrated with a box. The underlined sequence displays the domain with the WD-40 repeats in AtRRB1. The level of physicochemical amino acid conservation is shown below the alignment: full (*), high (:), and medium (.).

**Figure S2**

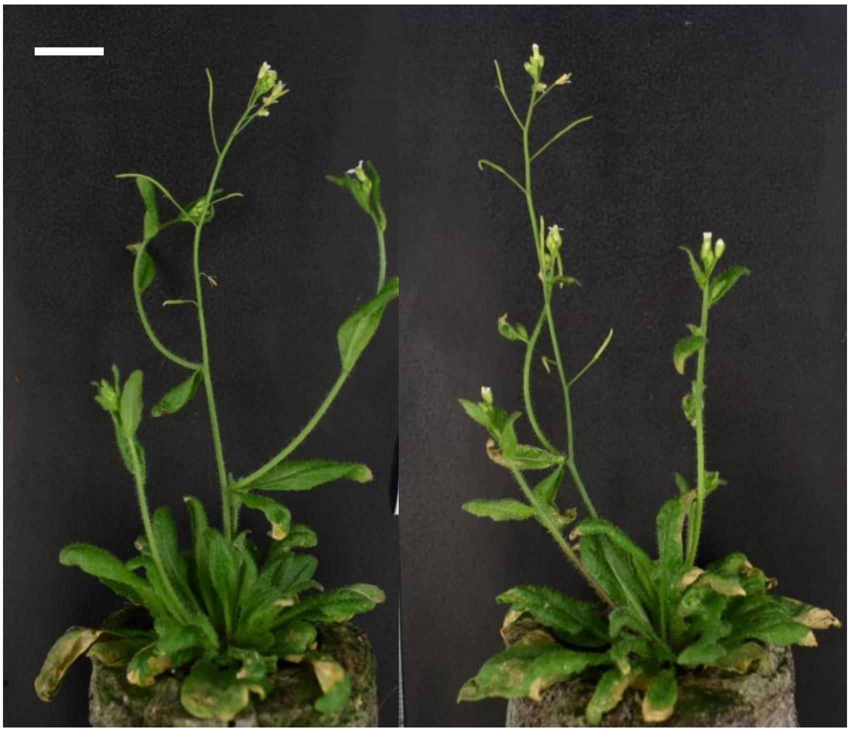

Figure S2. Morphology of Col-0 and *rrb1-1* Arabidopsis after 5 weeks of cultivation.
A. Col-0. B. *rrb1-1*. Size bar is 1 cm.

**Figure S3.**

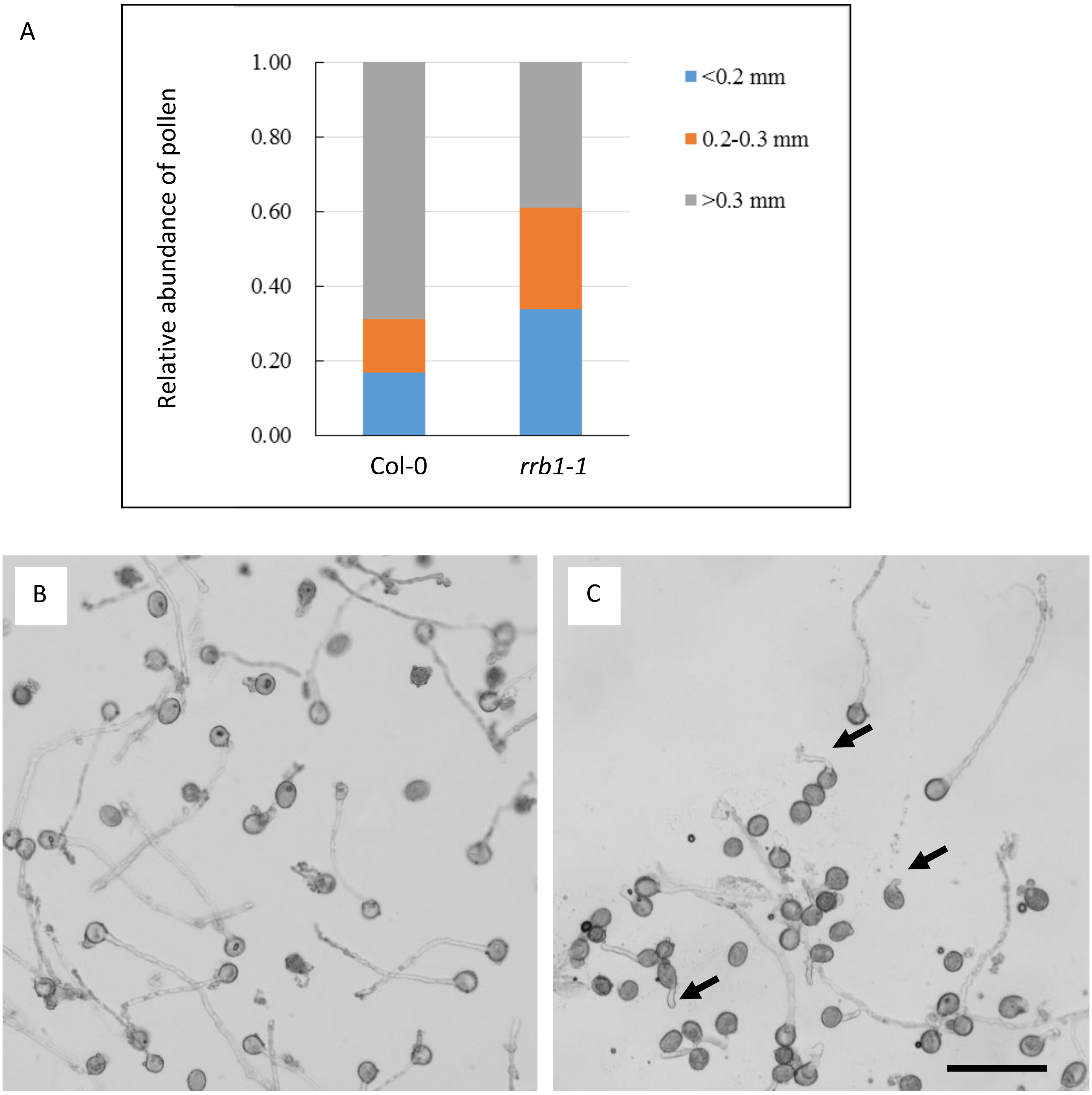


**Figure S3.** Pollen tube length is affected in *rrb1-1*.
A. Relative abundance of pollen from Col-0 and *rrb1*-1 with pollen tube length of <0.2 mm; 0.2-0.3 mm; and >0.3 mm (n>1000). B. Germinated pollen from Col-0. C. Germinated pollen from *rrb1-1*. Some pollen have long tubes like Col-0 and some pollen tubes are very short (arrows). Size bar is 50 $\mu$m.

**Supplementary tables**

**Table S1.** The T-DNA insertion GABI_837C04 in mutant rrb1-1 is embryo lethal.

| | **Total** | **Col-0** | **heterozygous** | **homozygous** |
|---|---|---|---|---|
| **N∘ plants** | 129 | 108 | 21 | 0 |
| **ratio** | | 83.7% | 16.3% | |

**Table S2.** Male and female transmission rates of the T-DNA insertion GABI_837C04.

| **Crossing** | **Tested** | **SulfaR** | **SulfaS** | **Transmission Rate** |
|---|---|---|---|---|
| Selfing | 400 | 64 | 336 | 16,0% |
| ♀Col-0 X ♂ *rrb1-1* | 207 | 34 | 173 | 16,4% |
| ♀ *rrb1-1* X ♂ Col-0 | 282 | 44 | 238 | 15,6% |
